# MeShClust^2^: Application of alignment-free identity scores in clustering long DNA sequences

**DOI:** 10.1101/451278

**Authors:** Benjamin T. James, Hani Z. Girgis

**Affiliations:** Bioinformatics Toolsmith Laboratory, Tandy School of Computer Science, The University of Tulsa, 800 South Tucker Drive, Tulsa, OK 74104, USA

## Abstract

Grouping sequences into similar clusters is an important part of sequence analysis. Widely used clustering tools sacrifice quality for speed. Previously, we developed MeShClust, which utilizes k-mer counts in an alignment-assisted classifier and the mean-shift algorithm for clustering DNA sequences. Although MeShClust outperformed related tools in terms of cluster quality, the alignment algorithm used for generating training data for the classifier was not scalable to longer sequences. In contrast, MeShClust^2^ generates semi-synthetic sequence pairs with known mutation rates, avoiding alignment algorithms. MeShClust^2^clustered 3600 bacterial genomes, providing a utility for clustering long sequences using identity scores for the first time.

## Introduction

Because of the rapid growth of sequenced genomes attributed to the lower cost in sequencing technologies, the bottleneck in unsolved problems has shifted from sequencing to sequence analysis^1^. Additionally, computer hardware improvements have been outpaced by this sequencing flood, leading to a call for efficient computational methods to sort and classify these data^2^.

Clustering DNA sequences is one of the main procedures applied in sequence analysis. Specialized sequence clustering tools have been developed for error reduction in reads produced by modern sequencing technologies^3–6^, redundancy detection^7^, finding representative sequences^8^, taxonomic profiling^9^, de-novo genome assembly^10–12^, and grouping expressed sequence tags^13^. Additionally, general purpose sequence clustering tools including MeShClust^14^, CD-HIT^5,15^, UCLUST^16^, DNACLUST^9^, d2_cluster^17^, mBKM^18^, and d2-vlmc^19^ have shown great applicability to many problems.

However, many of these methods — including CD-HIT, UCLUST, and DNACLUST — depend on a simple algorithm which lacks the ability to consistently produce optimal clusters when the exact value of the sequence identity score defining clusters is unknown. Recently, we developed MeShClust^14^ to address these concerns. MeShClust consists of these two components: (i) a self-supervised classifier for classifying sequence pairs into similar and dissimilar and (ii) the mean shift algorithm^20^ for clustering sequences.

The mean shift algorithm is a powerful general-purpose clustering algorithm which is able to find high quality clusters without beforehand knowing the number of clusters. Although wideley used in image processing^21–23^, the mean shift algorithm has rarely been applied in bioinformatics^24–26^.

Although it is the standard measure for sequence similarity, global alignment is such a computationally expensive algorithm. Its runtime is proportional to the square of the length of the sequences aligned, making alignment a bottleneck in most applications. This problem is especially visible in applications involving long sequences such as genomes or 3^*rd*^ generation sequencing reads. Additionally, most time in these clustering tools has been spent in calculating alignment, even with the numerous advances and heuristics in speeding up alignment. To minimize this alignment bottleneck, we utilized fast alignment-free k-mer statistics in MeShClust. In an instance of self-supervised learning, MeShClust computed alignment identity scores on sequence pairs found in the input data. Then, k-mer statistics were calculated on those same sequences so that combinations of these statistics could then be used to classify a sequence pair as above or below an identity score threshold. Using a combination of alignment-free k-mer statistics, MeShClust was able to compute high quality clusters, outperforming popular clustering tools.

However, MeShClust still had used alignment to generate labels for training the self-supervised classifier; this step required long time, especially on long sequences. Further, the alignment-free k-mer statistics used did not work well at lower identity values, as the statistics used were optimized for data sets above 60% identity. Therefore, MeShClust uses a global alignment algorithm directly, rather than the trained classifier, when the sequence identity threshold is below 60%. Furthermore, the classifier was trained on sequence pairs found in the input data. In some cases, there were too few sequence pairs with the needed identity scores for the classifier to be trained accurately or even trained at all. These three limitations have led us to develop MeShClust^2^.

We propose an alternative approach, which relies wholly on alignment-free methods using the foundation of MeShClust. In order to create identity scores to mimic alignment, we have formulated a procedure to simulate mutations as they occur in nature. Because the effect of these mutations is known, an identity score can be calculated to mimic the score produced by alignment. This means that the identity score can be calculated without actually calculating the alignment itself. We have applied this idea successfully in FASTCAR, which is a search tool for approximating the alignment identity score in linear time^27^. This alignment-free adaptation will allow for much faster clustering with comparable accuracy while extending the ability of MeShClust to cluster long DNA sequences. Clustering long DNA sequences is not currently possible to be done by any of the alignment-based (UCLUST, CD-HIT, and DNACLUST) or the alignment-assisted (MeShClust) clustering tools. We conclude that it is only possible for purely alignment-free algorithms to cluster this data. As a proof of concept, we applied MeShClust^2^ to clustering over 3600 bacterial genomes with average length of 3.4 mega base pairs.

## Methods

### Previous results

Recently we have developed a software tool called MeShClust^14^ for clustering DNA sequences. The tool depends on a classifier based on a self-supervised General Linear Model (GLM) and the mean shift algorithm, which is an instance of unsupervised learning algorithms. In supervised learning, manually-generated labeled examples are required for training, whereas labels are generated automatically in self-supervised learning. Other self-supervised systems were developed for detecting distal cis-regulatory modules^28^ and for annotating repeats^29^.

This classifier learns how to categorize sequence pairs as similar (with identity scores above a user-provided threshold) or dissimilar (with identity scores below a user-provided threshold). To provide labels for classification, the global alignment algorithm^30^ generates identity scores. These scores are then separated into the two aforementioned categories. Next, features based on k-mer statistics are calculated on each sequence pairs. The classifier learns a mapping from k-mer statistics to two labels: similar and dissimilar. In order to determine k-mer size, sequences are read in to gather their lengths. The k-mer length is then determined by the average length of the sequences. The k is determined by one less than the ceiling of the *log*_4_ of the average sequence length. The second part of MeShClust, the mean shift algorithm, is the backbone of the clustering algorithm for its non-parametric nature and ability to cluster data with high fidelity. Additionally, the mean shift algorithm allows for more accurate clustering of sequences than with simple, greedy approaches, which can often result in partial or incomplete clusters^14^.

Our previous approach has the following three drawbacks; (i) Alignment algorithms would take an impractically long time on long sequences, (ii) The classifier cannot be used reliably on low-identity sequences, and (iii) There may not be enough training sequence pairs with identities needed to train the classifier in the data set provided. To overcome these limitations, the alignment algorithm is eliminated by generating semi-synthetic sequence pairs with known mutation rates.

### Mutation models

As mentioned, we have completely removed alignment from MeShClust. To do so, we need a method which can yield alignment identity scores for training data. We utilized a generative idea to mutate real input sequences in order to produce a known identity score. Three mutation models are tested in our experiments. In most cases, some combination of some or all of these types of mutations should be used to accurately model natural mutations.

#### Single-point mutations

Single-point mutations are the most common mutations within gene coding regions, and are most of the mutations in high-identity alignments. These mutations are the insertion, deletion, or change of exactly one nucleotide at a spot. Multiple single-point mutations are possible and frequent. We ensured that these mutations do not overlap. In order to generate a mutated sequence with the same nucleotide composition as that of the real sequence, we select a nucleotide (A, C, G, or T) to be inserted or mismatched with nucleotide probabilities calculated on the real sequence. That is, an AT-rich region of DNA will have a high probability of inserting either an A or a T and a low probability of inserting a C or a G.

#### Block mutations

Block mutations (Figure 1) are common in higher mutation datasets. In these mutations, a group of nucleotides move together in a unit. These movements are classified into 3 different movements: insertion, deletion, and duplication. Insertion and deletion are block mutation versions of their single-point mutation counterparts. Similar to their single-point counterparts, nucleotide compositions are kept, and mutations do not overlap. However, it is more likely that a couple larger mutations occur rather than many single-point mutations. A certain number of mutations are allocated, and mutation lengths are randomly selected. The maximum block length must meet the conditions that (i) huge blocks (> 50 base pairs) are not considered, and (ii) at least 10 block mutations are used, negating the possibility that only a few block mutations are generated.

**Figure 1.**
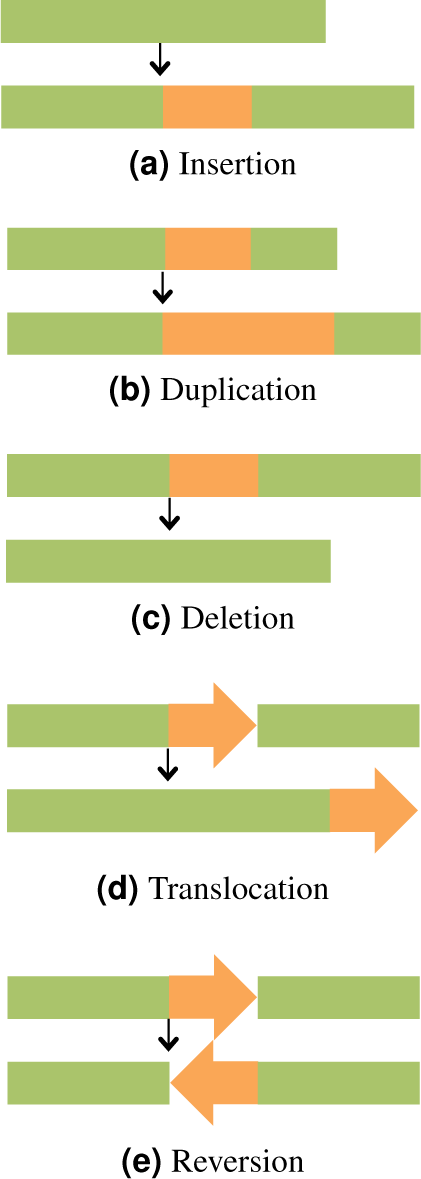
Non-single-point mutations — block mutations — including insertion, duplication, deletion, translocation, and reversion.

#### Infrequent mutations

Infrequent or uncommon mutations such as translocation and reversion can occur. See Figure 1. These mutation types are known to occur at the chromosomal level in entire genomes^31,32^. Translocation is the act of removing one segment and placing it somewhere else in the sequence. In reversion, a segment is reversed while occupying the same coordinates as the original segment. Alignment algorithms do not handle these mutations in a meaningful way. For example, a translocated segment is treated as a deleted block from one segment and as an inserted block in the other segment, extending the alignment length by double the length of the segment and lowering the identity score. However, the identity score should not be affected with translocation because identical nucleotides are still present in both sequences. For this reason, the generative method has the potential be more biologically relevant than alignment algorithms in handing translocation and reversion.

### Identity scores generation

In order to generate identity scores comparable to those due to the global alignment algorithm, the generative method mentioned above is utilized in MeShClust^2^. Simulating the process by which mutations are generated, we can make a feasible attempt to calculate identity scores. Knowing the effect of these mutations, we can calculate the number of matches and the alignment length. The ratio between these two numbers is the identity score. Table 1 shows how each mutation type affects the number of matches and the alignment length. Although these rules will not follow the actual alignment in all cases, when using conservative alignment parameters, this method yields accurate results. As mentioned, alignment algorithms do not handle the infrequent mutation types — translocation and reversion — in a biologically meaningful fashion. Therefore, it is not clear how each of them will affect the identity score. We assumed that a translocation even has the same effect as a matching event on the number of matches and the alignment length. However, we have not found a way to accurately model the effect of reversion on the alignment score. Handling these two mutation types requires further research. Because translocation and reversion are infrequent, the other types of mutations are sufficient to generate identity scores accurately.

**Table 1.** Here, the effects of various mutations on alignment identity score is shown. An identity score is the ratio between the number of matches and the alignment length. Initially, the number of matches and the alignment length are equal to the length of the original sequence. Each mutation type has a unique effect on these two numbers.

| Mutation type | Number of matches | Alignment length |
| --- | --- | --- |
| Insertion | No change | Increases by insertion length |
| Deletion | Decreases by deletion length | No change |
| Duplication | No change | Increases by duplication length |
| Mismatch | Decreases by 1 | No change |
| Translocation | No change | No change |
| Match | No change | No change |

### The self-supervised classifier

#### Generating balanced training and testing data sets

Previously in MeShClust, we used an alignment-free k-mer statistic, Czekanowski similarity^33^, to search the data set sequence pairs, upon which the alignment identity scores are computed. This has the potential to not have enough sequence pairs with identity scores above or below the user-specified threshold; if this is the case, the classifier cannot be trained. We have devised a more powerful approach. Instead of calculating alignments, each sequence which would have been chosen to be aligned with another is instead used for generating synthetic sequences with a desired range of identity scores. This range of identity scores not only spans equal amounts above and below the classification threshold but also generates alignments across all identity values. Here, the advantage of semi-synthetic mutated sequences is that they can be generated with almost any identity scores, producing a large number of sequence pairs enough to the train the classifier. Specifically, to generate the training and testing data sets, different lengths were uniformly sampled from the data provided to provide an accurate representation. Each sequence sampled would then be mutated 10 times, resulting in pairs with identity scores between 100% and 35%. Of these, 5 would be targeted into 5 equally-spaced bins above the identity threshold and the other 5 would be targeted into 5 equally spaced bins below. By default, 3000 training pairs and 3000 testing pairs are generated. This typically provides sufficient data to train the classifier, which is based on a General Linear Model (GLM).

#### The GLM-based classifier

GLMs are instances of supervised learning, and have applications in classification and regression tasks. In our past and current research, we have applied GLM to ranking protein structures^34–36^ and locating DNA microsatellites^37^. Using the semi-synthetic sequence pairs and their identity scores, we used various combinations of k-mer statistics^33^ as inputs to the GLM-based classifier. The GLM calculates weights for these k-mer statistics. Using the weighted combination of 3–5 k-mer statistics, the GLM-based classifier outputs a label predicting whether two sequences have an identity score above the user-provided threshold or not. Note that the classification is dependent on the data set and parameters chosen; therefore, the classifier can choose k-mer statistics to classify sequences more accurately. Next, we discuss how these 3–5 k-mer statistics are selected.

#### Selecting features

In the original version of MeShClust, a fixed feature set was used. That is, up to 4 k-mer feature pairs were used. These feature were: (i) length difference × Czekanowski, (ii) length difference^2^ × Manhattan^2^ (iii) Pearson coefficient, and (iv) length difference^2^ × Kulczynski2^2^. These features were optimized on few data sets with high (>60%) identity scores. Below 60% identity, we previously forced MeShClust to use the alignment algorithm directly. This was done since the GLM could not accurately classify sequences using these pre-selected features.

Due to the varying nature of data and identity values tested, we have now allowed many more k-mer feature combinations to be selected from in MeShClust^2^. In fact, we drew these combinations from a set of 9 single features which could be combined. These 9 features were gathered from our original study on alignment-free k-mer statistics^33^: (i) Euclidean, (ii) Manhattan, (iii) Czekanowski, (iv) Kulzynski2, (v) similarity ratio, (vi) normalized vectors, (vii) Pearson coefficient, (viii) earth mover distance, and (ix) length difference. These features/statistics represent a range of families that provide unique information and can be computed efficiently. Additionally, a large number of combinations between these features allows for flexibility. These combinations include multiplying these features together, such as Euclidean × length difference. Additionally, one or more of these 9 features can be squared, allowing for 162 possible combinations.

We applied an iterative, greedy approach to find the best combination of features. This approach is the same approach we used in FASTCAR^27^, which is a tool for approximating sequence identity scores in linear time. We utilized a training data set and a testing data set, both consisting of the semi-synthetic sequences mentioned earlier. The algorithm selects 3–5 features to allow for better information to be gathered at a reasonable time cost. At each step of the algorithm, each feature — not chosen yet — is added to the list of the selected features and the accuracy of the classifier is calculated. The feature combination which results in the highest accuracy becomes the list of the selected features. After three features are selected, a new feature is added to the list only if it leads to improving the accuracy. The algorithm stop if 5 features are chosen or no feature can improve the accuracy after selecting the first three features. This procedure results in a trained classifier, which is utilized by the mean shift algorithm as the distance kernel.

### The mean shift algorithm

We use the mean shift clustering algorithm due to its ability to form high quality clusters without specifying the number of clusters. Its bottom-up, iterative nature suits sequence clustering well and also provides well-chosen centers for the data. The mean shift algorithm takes the *mean* of similar sequences comprising one cluster, and then it *shift* s the center of the cluster to a better representation. Using a better estimate for the center of a cluster, more similar sequences can be found that accurately model the data. In this way, the data itself decides the shape of the clusters. As applied in MeShClust, the algorithm progresses in two stages, the accumulation stage and the updating stage.

The accumulation stage finds initial clusters from the data. Initially, the shortest sequence is chosen as the first cluster. Next, all sequences that the classifier finds close to the center are added to the cluster, and the mean shift is run on the new cluster, moving the center to a more accurate representation. This process is repeated until no new sequences are close to the center. At this point, the next closest sequence (according to Czekanowski similarity) is selected as the next cluster. Clusters are found and revised until no new sequences are left. This provides a semi-sorted list so that clusters are placed in a well ordered manner.

In the update stage, we take advantage of the placement of each cluster. Since clusters are placed in a sorted manner, neighboring clusters — the sequences in 5 clusters before and the 5 clusters after each cluster — are considered. The mean shift is then run on these similar sequences to form a new, more relevant center. Since the centers move, these centers may converge to a single center. Therefore, once the mean shift is run, if two centers are close to each other, then the clusters themselves are merged. This update process is performed until a fixed number of iterations is reached.

To show the effectiveness and speed of the generative model, we will demonstrate the ability of MeShClust^2^ to accurately cluster various data sets including entire bacterial genomes. Clustering long DNA sequences cannot be done using the alignment-based or the alignment-assisted clustering tools in practical time.

### Data sets

MeShClust was originally evaluated on a large microbial data set (Costello et al., 2009^38^), a set of viral genomes (viruSITE^39^), and multiple sets of synthetic sequences. The microbial data set includes about 1.1 million sequences with lengths around 200 base pairs (bp) to 400 bp sequenced from the 16S rRNA gene. Some of the synthetic data sets were formulated to generate sequences similar to those of the microbial (Costello) data set. These data sets were used in MeShClust^14^. The viral data set consists of 7 different families, although two families have two segments, leading to 9 total clusters. Sequences comprising a viral family have an average identity score of 40%–60%.

To showcase the true power of this alignment-free MeShClust^2^, we used a much larger data set consisting of bacterial genomes. All bacterial genomes from Ensembl Bacteria genomes (release 35) were downloaded, and all top-level FASTA sequences containing only one chromosome and no other contigs were considered in our evaluation^40^. This gave a subset of 3613 genomes from 43791 genomes. These genomes ranged in length from 112 kilo bp to over 14 mega bp, averaging 3.4 mega bp per genome. In order to provide a ground truth to this data, we generated three ground truth sets: families, genuses, and species. These ground truth sets were generated from Ensembl’s database, which included NCBI taxonomic identification. Using the NCBI Taxonomy Browser, each genome was assigned a family, genus, and species. For each ground truth set, all genomes were classified by family name, genus name, and species name for the three ground truths, respectively. This data can be accessed as Supplementary Data 1. A smaller subset including 92 genomes was used for faster tests. Similar to the larger data set, ground truth for this data set was generated at the family, genus, and species level.

### Related tools

We compared MeShClust^2^ to popular clustering algorithms such as CD-HIT, DNACLUST, and UCLUST as well as the alignment-assisted MeShClust on the aforementioned data sets. All of these tools were run with default parameters except for thread-number parameters and sequence identity parameters to provide an equal comparison. All tools were run on a Dell Precision Tower 5210 with 32 GB RAM and 20x Intel Xeon E5-2630 v4 running GNU/Linux (Ubuntu 17.10). MeShClust^2^ source code is available as Supplementary Data 2 and on GitHub t89(https://github.com/TulsaBioinformaticsToolsmith/MeShClust2).

## Results

### Evaluation measures

In order to determine the quality of clusters produced by the aforementioned tools, we utilized the following seven different evaluation measures: (i) intra-cluster similarity, (ii) inter-cluster similarity, (iii) silhouette score, (iv) purity, (v) normalized mutual information (NMI), (vi) time requirement, and (vii) memory usage. Of the first five evaluation measures, which are cluster quality measures, the first three are independent of the test type, but purity and NMI require a ground truth, i.e. known clusters.

#### General evaluation measures

The first three measures, intra-cluster similarity, inter-cluster similarity, and silhouette^41^, use alignment to grade the clusters. Intra-cluster similarity measures the average alignment identity between each sequence and the center of its cluster. Inter-cluster similarity measures the average alignment identity among a center of a cluster and the centers of other clusters. Intra-cluster and inter-cluster distance can provide insight into how the predicted clusters look, as a high intra-cluster similarity and a low inter-cluster similarity can provide bounds on how similar sequences in each cluster are and how separated the clusters are.

Silhouette measure^41^ combines the two previous measures. As seen by Equation (1), silhouette measure is a good measure for balancing between how close sequences are and how far away the clusters are.

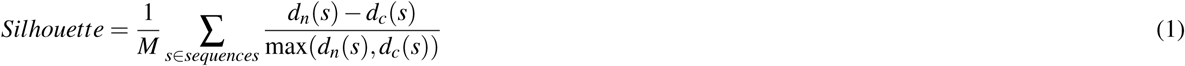

Here, *d_c_*(*s*) is the distance between the sequence *s* and the sequence at the center of *s*’s own cluster, and *d_n_*(*s*) is the distance between s and the center of the next closest “neighboring” cluster. The distance is then obtained by subtracting the identity score from 100%. In this equation, *M* is the number of sequences in the data set. Because the denominator is normalized to the maximum of the two distances, the output is scaled between 1 and-1. A high score indicates that sequences in a cluster are tighter and clusters are far apart from each other; a low score can indicate high proximity among clusters or general poor clustering. In this way, it provides a single score measuring the quality of clusters produced without knowing the ground truth. Here, the silhouette measure only evaluates clusters with at least 5 members. Otherwise, artificially inflated numbers can appear by clustering small clusters, especially those clusters consisting of one sequence. The silhouette value for that sequence would be 1, as all cluster centers get a silhouette value of 1. Unfortunately, this measure cannot tell whether the data actually matches the ground truth, but is the best estimation when ground truth is not available.

#### Evaluation measures utilizing the ground truth

But, on data where there is a ground truth, traditional clustering evaluation measures can be used. Examples of these are purity (Equation 2)^42^ and NMI (Equation 5)^42^. These measures compare all clusters found,*F*, to the actual clusters, *A*, in a set of *N* sequences. Purity is the measure of how “mixed” or “pure” a cluster is, if there are clusters with sequences from multiple actual clusters, that cluster will probably have a lower purity score.

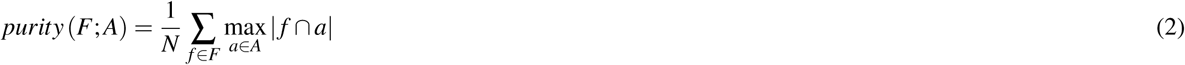

NMI, coming from the world of information theory, considers the probability that *F* and *A* contain the same “mututal” information. In order to normalize this, the mutual information is divided by the average entropy of *F* and *A* individually. Equations (3), (4), and (5) describe mutual information, entropy, and NMI, respectively.

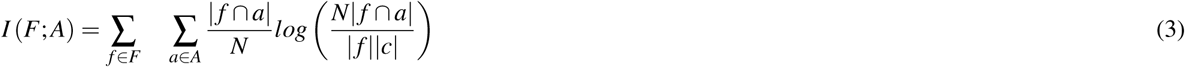

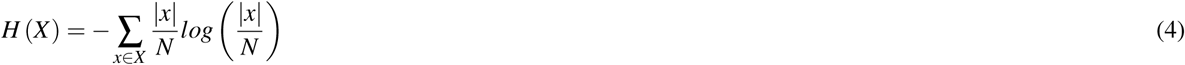

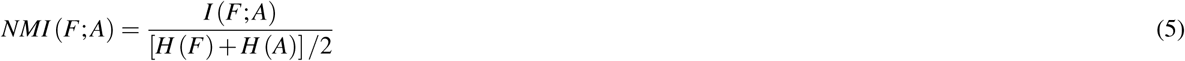

On the bacterial data sets, because we have hierarchical ground truth files — family, genus, and species — we attempt to gauge the performance across all three. We wish to produce a single number from the predicted clusters to show their overall similarity to the three ground truth sets. To do so, we first combine the NMI and the purity on each ground truth set using the geometric mean. Then, we take the average of these geometric means to compute the total score (Equation 6).

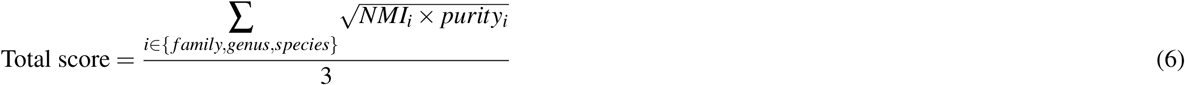

With these evaluation criteria, we move on to the evaluations on the data sets mentioned.

### Evaluation on the small subset of the bacterial genomes using three the mutation models

Other tools such as the alignment-assisted MeShClust, UCLUST and CD-HIT cannot cluster genomes of this size. Long sequences are a weakness in term of speed of global alignment algorithms, even with modern time-cutting heuristics. Therefore, this data set can only be clustered by MeShClust^2^. Since different mutation models were conceived to model biological mutations, we compared all of these models on the small subset of bacterial genomes. Specifically, we considered three mutation models. The first model is based on single-point mutations only; we call this model “Single.” The second model is based on single-point mutations as well as block mutations without translocations; we call this model “Both.” The third model is the same as the second, but it allows for translocations; we call this model “Both+Trans.”

As seen in the results of this experiment (Table 2), the Single model does not provide the same quality of clusters that are provided by the Both model or the Both+Trans model. Additionally, the single mutation model on lower identity values takes much more time because the mutation model requires no overlaps. On long sequences, this search can take a lot of time.

**Table 2.**
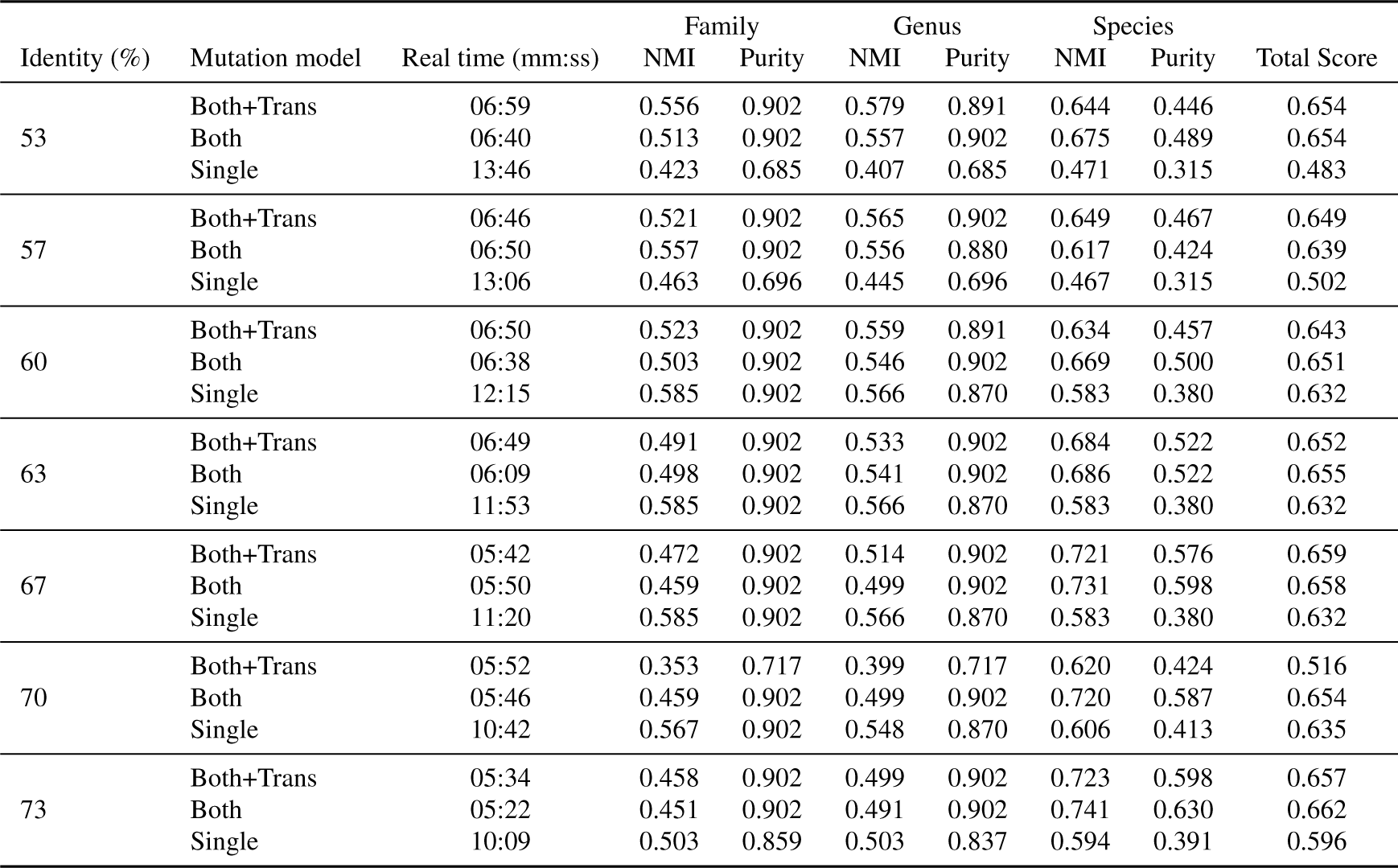
Performance of MeShClust^2^ on a small subset of the bacterial genomes using three ground truth sets representing families, genuses, and species. The purpose of this experiment is to determine which mutation model is most suitable to clustering full genomes. We compared the performances due to the following three mutation models: (i) single point: “Single”; (ii) single point and block without translocation: “Both”, and (iii) single point and block with translocation: “Both+Trans.” This experiment was conducted using 7 identity thresholds. The NMI (Equation 5) is a measure of the intersection between the predicted clusters and the real ones — the higher, the better. The purity (Equation 2) is a measure of how pure or how mixed the predicted clusters — the higher, the better. The total score (Equation 6) is a composite score of the NMI and the purity on the family, genus, and species levels — the higher, the better.

| Identity (%) | Mutation model | Real time (mm:ss) | Family |  | Genus |  | Species |  | Total Score |
| --- | --- | --- | --- | --- | --- | --- | --- | --- | --- |
|  |  |  | NMI | Purity | NMI | Purity | NMI | Purity |  |
| 53 | Both+Trans | 06:59 | 0.556 | 0.902 | 0.579 | 0.891 | 0.644 | 0.446 | 0.654 |
|  | Both | 06:40 | 0.513 | 0.902 | 0.557 | 0.902 | 0.675 | 0.489 | 0.654 |
|  | Single | 13:46 | 0.423 | 0.685 | 0.407 | 0.685 | 0.471 | 0.315 | 0.483 |
| 57 | Both+Trans | 06:46 | 0.521 | 0.902 | 0.565 | 0.902 | 0.649 | 0.467 | 0.649 |
|  | Both | 06:50 | 0.557 | 0.902 | 0.556 | 0.880 | 0.617 | 0.424 | 0.639 |
|  | Single | 13:06 | 0.463 | 0.696 | 0.445 | 0.696 | 0.467 | 0.315 | 0.502 |
| 60 | Both+Trans | 06:50 | 0.523 | 0.902 | 0.559 | 0.891 | 0.634 | 0.457 | 0.643 |
|  | Both | 06:38 | 0.503 | 0.902 | 0.546 | 0.902 | 0.669 | 0.500 | 0.651 |
|  | Single | 12:15 | 0.585 | 0.902 | 0.566 | 0.870 | 0.583 | 0.380 | 0.632 |
| 63 | Both+Trans | 06:49 | 0.491 | 0.902 | 0.533 | 0.902 | 0.684 | 0.522 | 0.652 |
|  | Both | 06:09 | 0.498 | 0.902 | 0.541 | 0.902 | 0.686 | 0.522 | 0.655 |
|  | Single | 11:53 | 0.585 | 0.902 | 0.566 | 0.870 | 0.583 | 0.380 | 0.632 |
| 67 | Both+Trans | 05:42 | 0.472 | 0.902 | 0.514 | 0.902 | 0.721 | 0.576 | 0.659 |
|  | Both | 05:50 | 0.459 | 0.902 | 0.499 | 0.902 | 0.731 | 0.598 | 0.658 |
|  | Single | 11:20 | 0.585 | 0.902 | 0.566 | 0.870 | 0.583 | 0.380 | 0.632 |
| 70 | Both+Trans | 05:52 | 0.353 | 0.717 | 0.399 | 0.717 | 0.620 | 0.424 | 0.516 |
|  | Both | 05:46 | 0.459 | 0.902 | 0.499 | 0.902 | 0.720 | 0.587 | 0.654 |
|  | Single | 10:42 | 0.567 | 0.902 | 0.548 | 0.870 | 0.606 | 0.413 | 0.635 |
| 73 | Both+Trans | 05:34 | 0.458 | 0.902 | 0.499 | 0.902 | 0.723 | 0.598 | 0.657 |
|  | Both | 05:22 | 0.451 | 0.902 | 0.491 | 0.902 | 0.741 | 0.630 | 0.662 |
|  | Single | 10:09 | 0.503 | 0.859 | 0.503 | 0.837 | 0.594 | 0.391 | 0.596 |

Recall that the total score metric (Equation 6) is an attempt to produce a single number to gauge the performance on all ground truths — families, genuses, and species. Averaging over the 7 identities, the average total score for the Single model was 0.588, for the Both model was 0.654, and for the Both+Trans was 0.633. This can be explained by the fact that genomes often have long segments missing/inserted in reference to each other. These missing/inserted longer segments are best explained by block mutations. We observed that there is no added value due to translocation. Therefore, the mutation model combining single-point mutations and block mutations without translocation is the most suitable mutation model to the bacterial genomes.

### Evaluation on the small subset of the bacterial genomes using a shorter k-mer

One of our goal is to make our tool work on a regular personal computer — no need for a supercomputer. To this end, a shorter k-mer is needed to reduce the memory requirement to what is available on a personal computer. Reduced k-mer values are often used to reduce physical memory constraints by hardware, since the memory required is proportional to 4^*k*^. However, we ensured that using a (k-1)-mer produces comparable results to the results due to the default k-mer. Next, we compare the performance of MeShClust^2^ using the default 10-mer to the performance due to 9-mer on the same 92 bacterial genomes. We used the best performing mutation model: Both — both single-point and block mutations — only, allowing us to focus on the effects of the reduced k. Table 3 shows the results. There is not much of a drop-off in terms of quality between using 10 or 9. In fact, the average total score due to 9-mer is 0.644 and the average total score due to 10-mer is 0.654. These numbers indicate that using 9-mer should result in comparable results obtained due to the default (10-mer). In the next experiment, using 9-mer allowed us to cluster about 3600 bacterial genomes on a computer with 32 GB of RAM.

**Table 3.** Comparing the performance of MeShClust^2^ using the default based on 10-mer to the performance based on 9-mer. Using 9-mer requires about 25% of the memory required if 10-mer is used. This way, clustering a large number of bacterial genomes can be done on a desktop without the need of a supercomputer or specialized hardware. The NMI (Equation 5) is a measure of the intersection between the predicted clusters and the real ones — the higher, the better. The purity (Equation 2) is a measure of how pure or how mixed the predicted clusters — the higher, the better. The total score (Equation 6) is a composite score of the NMI and the purity on the family, the genus, and the species levels — the higher, the better.

| Identity (%) | k-mer | Real time | Max memory (MB) | Family |  | Genus |  | Species |  | Total Score |
| --- | --- | --- | --- | --- | --- | --- | --- | --- | --- | --- |
|  |  |  |  | NMI | Purity | NMI | Purity | NMI | Purity |  |
| 53 | 9 | 6:34.14 | 6002 | 0.579 | 0.902 | 0.560 | 0.870 | 0.601 | 0.413 | 0.639 |
|  | 10 | 6:39.89 | 10923 | 0.513 | 0.902 | 0.557 | 0.902 | 0.675 | 0.489 | 0.654 |
| 57 | 9 | 6:29.34 | 6249 | 0.539 | 0.891 | 0.521 | 0.859 | 0.619 | 0.424 | 0.624 |
|  | 10 | 6:49.69 | 12382 | 0.557 | 0.902 | 0.556 | 0.880 | 0.617 | 0.424 | 0.639 |
| 60 | 9 | 6:10.48 | 6466 | 0.464 | 0.837 | 0.513 | 0.837 | 0.654 | 0.457 | 0.608 |
|  | 10 | 6:38.17 | 12972 | 0.503 | 0.902 | 0.546 | 0.902 | 0.669 | 0.500 | 0.651 |
| 63 | 9 | 5:53.06 | 6859 | 0.507 | 0.902 | 0.551 | 0.902 | 0.678 | 0.500 | 0.654 |
|  | 10 | 6:08.91 | 13659 | 0.498 | 0.902 | 0.541 | 0.902 | 0.686 | 0.522 | 0.655 |
| 67 | 9 | 5:28.08 | 6695 | 0.509 | 0.902 | 0.552 | 0.902 | 0.686 | 0.511 | 0.658 |
|  | 10 | 5:50.32 | 14210 | 0.459 | 0.902 | 0.499 | 0.902 | 0.731 | 0.598 | 0.658 |
| 70 | 9 | 5:19.57 | 6838 | 0.484 | 0.902 | 0.526 | 0.902 | 0.718 | 0.565 | 0.662 |
|  | 10 | 5:45.71 | 14534 | 0.459 | 0.902 | 0.499 | 0.902 | 0.720 | 0.587 | 0.654 |
| 73 | 9 | 5:11.16 | 6867 | 0.466 | 0.902 | 0.507 | 0.902 | 0.729 | 0.598 | 0.661 |
|  | 10 | 5:22.37 | 14840 | 0.451 | 0.902 | 0.491 | 0.902 | 0.741 | 0.630 | 0.662 |

### Evaluation on the entire bacterial data set

Using the results from the previous experiment (Table 3), there is a little difference between the default k-mer size and one less than that. As the whole bacterial data set would not fit in the memory of the testing machine, the reduced k-mer size requires less memory. Next, we evaluated MeShClust^2^ on the entire bacterial data set, which consists of of 3613 bacterial genomes. These results suggest that an identity score of 70% produces clusters that represent bacterial families (NMI: 0.831, Purity: 0.918); and an identity score of 80% produces clusters representing genuses (NMI: 0.888, Purity: 0.967). When the predicted clusters are compared to the species clusters, the NMI and the purity values increase as the identity threshold increases — as expected. These results are in line with the family-genus-species hierarchy.

Next, we also compared the ground truths against each other as benchmarks. Evaluating them against each other shows the effect of good/bad clustering on the evaluation measures, attaching a number to a known quantity; it also shows the biological relevance of predicted clusters. The information encoded in these clusters is shown in the NMI, and we can see the effect of different taxa on this measure. Similarly, as the purity measure encodes the cohesiveness and commonality of the clusters, the effect is seen. We can see in many cases, the difference between adjacent taxa (e.g. family and genus) is similar to MeShClust^2^clusters across multiple identity values. On most identity values, MeShClust^2^ produces clusters with NMI between species and genus when evaluated on family. This implies that MeShClust produces clusters with similar information to a cluster hierarchy between species and genus. On genus, many of the identities around 73% have higher NMI than species against genus and higher purity against family. This shows that most of the clusters are clustered in that range. On species, above 67%, both the NMI and Purity are better than genus or family evaluated on species ground truth. In sum, MeShClust does produce biologically-relevant clusters.

**Table 4.** The performance of MeShClust^2^ on the 3613 bacterial genomes. We tried several identity thresholds and evaluated the performance on true clusters representing bacterial families, genuses, and species. The NMI measures the overlap between the predicted clusters and the true clusters. The Purity measures the composition purity of the predicted clusters; it is preferable that elements comprising a predicted cluster to belong to only one true cluster. In the last three rows of the table, we used the true clusters as if they were the predicted clusters. For example, the row labeled “Family” shows the NMI and the Purity of the families, the genuses, and the species on the family clusters (treating families as the predicted clusters). We did so to provide benchmarks on the NMI and the purity values.

| Identity (%) | Real time (mm:ss) | Max memory (MB) | Family |  | Genus |  | Species |  |
| --- | --- | --- | --- | --- | --- | --- | --- | --- |
|  |  |  | NMI | Purity | NMI | Purity | NMI | Purity |
| 53 | 36:18 | 29328 | 0.704 | 0.603 | 0.738 | 0.537 | 0.667 | 0.353 |
| 57 | 36:44 | 29397 | 0.754 | 0.706 | 0.799 | 0.650 | 0.728 | 0.423 |
| 60 | 37:11 | 29672 | 0.804 | 0.777 | 0.839 | 0.708 | 0.754 | 0.466 |
| 63 | 36:33 | 29663 | 0.786 | 0.750 | 0.825 | 0.683 | 0.748 | 0.453 |
| 67 | 39:12 | 29755 | 0.822 | 0.890 | 0.874 | 0.837 | 0.818 | 0.589 |
| 70 | 36:11 | 29895 | 0.831 | 0.918 | 0.886 | 0.872 | 0.828 | 0.624 |
| 73 | 35:23 | 29920 | 0.826 | 0.930 | 0.887 | 0.893 | 0.833 | 0.647 |
| 77 | 37:27 | 29730 | 0.794 | 0.912 | 0.860 | 0.882 | 0.821 | 0.767 |
| 80 | 34:23 | 30278 | 0.817 | 0.992 | 0.888 | 0.967 | 0.847 | 0.854 |
| 83 | 34:28 | 28266 | 0.814 | 0.990 | 0.883 | 0.960 | 0.849 | 0.856 |
| Family | – | – | 1.000 | 1.000 | 0.908 | 0.743 | 0.734 | 0.364 |
| Genus | – | – | 0.907 | 0.980 | 1.000 | 1.000 | 0.802 | 0.518 |
| Species | – | – | 0.739 | 0.981 | 0.805 | 0.995 | 1.000 | 1.000 |

Clustering these long genomes (3.4 mega bp long on average) took MeShClust^2^ only 34–39 minutes. Again, no other clustering tool can use identity thresholds to cluster these 3600 bacterial genomes. These results demonstrate the success of using semi-synthetic sequence pairs with known mutation rates as efficient replacement to identities due to alignment algorithms.

### Evaluation on viral genomes

Recall that the classifier of the original MeShClust was trained on labels generated by the global alignment algorithm. This classifier is used to predict whether or not the similarity between two sequences are greater than the provided threshold. If the identity threshold is less 60%, MeShClust uses the global alignment algorithm instead of the alignment-assisted classifier because the predetermined features were not informative enough to distinguish sequences at this low identity. Additionally, given the small size of this data set, there were not enough sequences to generate a balanced training set. By balanced, we mean that the training set should include roughly equal number of sequence pairs with identity scores above the threshold and pairs with scores below the threshold. The classifier of MeShClust^2^ remedies these two limitations. First, the new classifier utilizes a greedy algorithm for selecting features among a much larger number of features, whereas MeShClust’s classifier uses up to four predetermined features. Second, because MeShClust^2^ generates semi-synthetic sequence pairs for training and testing, it can generate a large enough number sufficient to train the classifier. With these two ideas, MeShClust^2^ is able to cluster sequences at low-identity thresholds utilizing only alignment-free methods.

Table 5 show the performances of UCLUST, MeShClust, and MeShClust^2^ on 96 viral genomes. The alignment-based MeShClust still leading in terms of the NMI and the purity, overall. UCLUST and the alignment-free MeShClust^2^ obtained comparable NMI and purity scores, overall. However, MeShClust^2^ is much faster than both of MeShClust and UCLUST. Yet, the alignment-free MeShClust^2^ has not outperformed the alignment-based MeShClust. However, this task — clustering viral genomes at low identities — was not possible using alignment-free methods prior to MeShClust^2^. Therefore, the novel ideas implemented in MeShClust^2^ represent true progress in the alignment-free field.

**Table 5.** Evaluations on the viral data set, which consists of 9 clusters. Only UCLUST, MeShClust, and MeShClust^2^ can cluster sequences with low sequence identities. The NMI is a measure of the overlap between the predicted clusters and the true clusters. The purity is a measure of how mixed or how pure the predicted clusters are; these clusters are desired to include sequences that belong to one true cluster. The last column shows the geometric mean of the NMI and the purity, combining them into one score. An error in the evaluation script, assuming all sequences were in UCLUST’s output, has been fixed; this may lead to slightly different purity and NMI values for UCLUST from those reported in our earlier study^14^.

| Identity (%) | Tool | Real time (m:ss) | Max memory (MB) | NMI | Purity | $\sqrt{NMI \times Purity}$ |
| --- | --- | --- | --- | --- | --- | --- |
| 43 | UCLUST | 0:12.79 | 72 | 0.064 | 0.438 | 0.000 |
|  | MeShClust | 0:13.10 | 87 | 0.583 | 0.573 | 0.577 |
|  | MeShClust<sup>2</sup> | 0:00.55 | 29 | 0.000 | 0.438 | 0.000 |
| 47 | UCLUST | 0:22.21 | 74 | 0.502 | 0.656 | 0.573 |
|  | MeShClust | 0:14.45 | 88 | 0.677 | 0.667 | 0.671 |
|  | MeShClust<sup>2</sup> | 0:00.80 | 31 | 0.710 | 0.823 | 0.764 |
| 50 | UCLUST | 0:35.92 | 76 | 0.735 | 0.938 | 0.830 |
|  | MeShClust | 0:40.02 | 87 | 0.865 | 0.917 | 0.890 |
|  | MeShClust<sup>2</sup> | 0:01.15 | 32 | 0.665 | 0.771 | 0.716 |
| 53 | UCLUST | 0:40.61 | 77 | 0.604 | 0.938 | 0.752 |
|  | MeShClust | 1:13.17 | 87 | 0.866 | 1.000 | 0.930 |
|  | MeShClust<sup>2</sup> | 0:01.41 | 34 | 0.688 | 0.938 | 0.803 |
| 57 | UCLUST | 0:41.80 | 78 | 0.584 | 0.938 | 0.740 |
|  | MeShClust | 2:38.61 | 87 | 0.687 | 1.000 | 0.828 |
|  | MeShClust<sup>2</sup> | 0:02.03 | 38 | 0.670 | 0.875 | 0.765 |

### Evaluation on the microbial data set

Finally, we want to compare the alignment-free MeShClust^2^ to the same tools on the same data sets used in the original evaluation of MeShClust^14^. The microbial data set was clustered using these five identity thresholds: 83%, 87%, 90%, 93%, and 97%. As seen in Table 6, again MeShClust^2^ outperforms the other tools in terms of Silhouette scores on every single identity, and does so by a large margin. These scores are similar to those due to the alignment-assisted MeShClust. The alignment-free MeShClust^2^ and the alignment-assisted MeShClust have very comparable average Silhouette score (0.462 vs. 0.456). As expected, the time required has drastically reduced by 25%–45% (now comparable to the other tools). Most of this speed-up is attributed to the reduction in time required for training the self-supervised GLM-based classifier, which had previously been trained using identity scores calculated by the quadratic alignment algorithm. Further, the alignment-free MeShClust^2^ had lower memory requirement by 16%–22%. These results demonstrate that that the alignment-free identity scores can replace the ones calculated by the quadratic alignment algorithm successfully, leading to reduction in both time and memory.

**Table 6.** Comparison on the microbial data set, which includes about 1.1 million sequences. The Silhouette score measures how tight and how separable the predicted clusters are — the higher, the better. The Intra-cluster similarity measures how similar sequences of one predicted cluster to each other — the higher, the better. The Inter-cluster similarity measures how similar the centers of different predicted clusters to each other — the lower, the better.

| Identity (%) | Tool | Silhouette | Intra-cluster (%) | Inter-cluster (%) | Real time (m:ss) | Max memory (MB) |
| --- | --- | --- | --- | --- | --- | --- |
| 83 | CD-HIT | 0.186 | 81.366 | 62.893 | 1:52 | 973 |
|  | DNACLUST | 0.169 | 82.006 | 63.163 | 1:06 | 2367 |
|  | MeShClust<sup>2</sup> | 0.448 | 85.742 | 63.030 | 3:33 | 871 |
|  | MeShClust | 0.450 | 89.074 | 65.034 | 5:52 | 1038 |
|  | UCLUST | 0.299 | 85.951 | 63.651 | 2:43 | 412 |
| 87 | CD-HIT | 0.227 | 85.430 | 63.977 | 1:43 | 976 |
|  | DNACLUST | 0.218 | 86.102 | 64.214 | 1:40 | 2361 |
|  | MeShClust<sup>2</sup> | 0.425 | 88.468 | 63.771 | 2:58 | 834 |
|  | MeShClust | 0.468 | 90.455 | 65.751 | 5:03 | 1051 |
|  | UCLUST | 0.310 | 88.611 | 64.469 | 3:49 | 412 |
| 90 | CD-HIT | 0.230 | 87.850 | 64.620 | 1:52 | 978 |
|  | DNACLUST | 0.197 | 88.395 | 64.936 | 2:03 | 2359 |
|  | MeShClust<sup>2</sup> | 0.460 | 91.165 | 64.658 | 3:44 | 834 |
|  | MeShClust | 0.446 | 91.718 | 65.154 | 4:57 | 1071 |
|  | UCLUST | 0.302 | 90.672 | 65.308 | 1:14 | 412 |
| 93 | CD-HIT | 0.217 | 90.337 | 65.446 | 1:07 | 982 |
|  | DNACLUST | 0.194 | 91.155 | 65.828 | 2:30 | 2369 |
|  | MeShClust<sup>2</sup> | 0.504 | 92.852 | 65.151 | 3:41 | 844 |
|  | MeShClust | 0.454 | 92.998 | 65.629 | 5:47 | 1083 |
|  | UCLUST | 0.246 | 92.612 | 66.343 | 1:41 | 412 |
| 97 | CD-HIT | 0.179 | 94.327 | 66.785 | 1:28 | 993 |
|  | DNACLUST | 0.156 | 94.958 | 67.185 | 2:16 | 2365 |
|  | MeShClust<sup>2</sup> | 0.477 | 94.248 | 66.721 | 2:57 | 871 |
|  | MeShClust | 0.464 | 95.321 | 66.553 | 5:20 | 1102 |
|  | UCLUST | 0.073 | 95.905 | 67.304 | 3:10 | 412 |

### Evaluation on synthetic data sets

We evaluated MeShClust^2^, the alignment-assisted MeShClust and three related tools on synthetic data sets. These data sets were generated using different mutation rates. Results are available as Supplementary Data 3. MeShClust^2^ produced stable clusters — similar to those produced by the alignment-assisted MeShClust — and outperformed the three related tools. These results demonstrate the success of the alignment-free MeShClust^2^.

## Conclusions

Although highly popular, UCLUST and CD-HIT utilize simple, greedy algorithms which trade quality for speed. Additionally, they depend on global alignment, a costly algorithm whose run time increases quadratically with regard to sequence length. While not an issue on gene-length sequences, long sequences present a challenge for alignment algorithms. Alternatively, tens of alignment-free k-mer statistics have been proposed. However, their scores are not as biologically relevant as the identity scores produced by alignment algorithms.

To overcome these limitations, we developed MeShClust, which (i) utilizes a classifier that maps efficient k-mer statistics to a classification according to biologically-meaningful identity scores, and (ii) implements the mean shift algorithm to produce quality clusters. This combination allows for the quality of the mean shift to shine with the speed of k-mer statistics.

However, this application has some drawbacks — with the classifier in particular. Firstly, MeShClust was not completely alignment-free, as identity scores are necessary to train the classifier. Secondly, the features gathered may not be flexible, specifically because the fixed feature set was optimized for high identity gene data sets. Finally, MeShClust depended on a high number of sequence pairs to be aligned in order to generate training data for the classifier.

We have addressed these concerns with a generative and adaptive approach. Using this approach, semi-synthetic sequences are generated with known mutation rates; therefore, the identity scores can be calculated without using the alignment algorithm. Because synthetic data is generated, a variety of identity values and a plethora of training data can be generated with ease. Since this data can vary, we developed a classifier training method, which can adapt to these synthetic training sets. These adaptations have allowed clustering on large data sets much faster than previous approaches.

As a proof of concept, a data set of over 3600 genomes has been able to be clustered using this method. We have shown that with speed not possible with alignment, these clusters still have biological relevance that alignment would provide. Finally, a user-friendly identity parameter wraps this method, allowing these large sequences to be clustered similarly to alignment.

## Data availability

The C++ source code of MeShClust^2^ available on GitHub t89 (https://github.com/TulsaBioinformaticsToolsmith/MeShClust2) and as Supplementary Data 1.

## Acknowledgements

The authors would like to thank Alexander Baumgartner for his help with data processing and coding the module for generating mutated sequences. This research was supported mainly by funds from the Oklahoma Center for the Advancement of Science and Technology [PS17-015] and in part by internal funds provided by the College of Engineering and Natural Sciences and the Tulsa Undergraduate Research Challenge (TURC) Program at the University of Tulsa.

## Author contributions statement

H.Z.G. conceived the idea and designed the experiments. B.T.J. developed the software, conducted the experiments, and produced the results. Both authors wrote and reviewed the manuscript.

